# Spatio-Temporal Pattern of Appraising Social and Emotional Relevance: Evidence from Event-Related Brain Potentials

**DOI:** 10.1101/230961

**Authors:** Annekathrin Schacht, Pascal Vrtička

## Abstract

Social information is highly intrinsically relevant for the human species because of its direct link to guiding physiological responses and behavior. Accordingly, extant functional magnetic resonance imaging (fMRI) data suggest that social content may form a unique stimulus dimension. It remains largely unknown, however, how neural activity underlying social (versus nonsocial) information processing temporally unfolds, and how such social information appraisal may interact with the processing of other stimulus characteristics, particularly emotional meaning. Here, we presented complex visual scenes differing in both social (versus nonsocial) and emotional relevance (positive, negative, neutral) intermixed with scrambled versions of these pictures to N= 24 healthy young adults. Event-related brain potentials (ERPs) to intact pictures were examined for gaining insight to the dynamics of appraisal of both dimensions, implemented within the brain. Our main finding is an early interaction between social and emotional relevance due to enhanced amplitudes of early ERP components to emotionally positive pictures of social compared to nonsocial content, presumably reflecting rapid allocation of attention and counteracting an overall negativity bias. Importantly, our ERP data show high similarity with previously observed fMRI data using the same stimuli, and source estimations located the ERP effects in overlapping occipito-temporal brain areas. Our new data suggest that relevance detection may occur already as early as around 100 ms after stimulus onset and may combine relevance checks not only examining intrinsic pleasantness/emotional valence, but also social content as a unique, highly relevant stimulus dimension.

## Introduction

Humans are highly social beings (Aronson, 1980; Tomasello, 2014). Hence, social information is assumed to be of high intrinsic relevance to humans due to its direct link to guiding physiological responses and behavior (Hariri et al., 2002; Keltner and Kring, 1998). A prominent evolutionary theory, the *social brain hypothesis*, even postulates that primates – including humans – have evolved unusually large brains for body size compared to all other vertebrates as a means to manage their unusually complex social systems (Dunbar, 1998, 2009). During the last two and a half decades, much research has thus been dedicated to better understand the functioning of the so-called social brain in a newly emerging field termed *social cognitive affective neuroscience* (Adolphs, 2003; Cacioppo and Berntson, 1992; Lieberman, 2007). Along these lines, social stimuli are argued to constitute the most emotionally evocative stimuli for humans, providing vital clues for survival throughout the life span by promoting both affiliative (e.g. attachment, reproduction) as well as protective (e.g. vigilance toward threatening encounters, protection of territory and significant others) behaviors (Insel, 2010; Norris et al., 2004; Porges, 2003). Accordingly, social interactions are thought to be motivated by emotions directing long-term social goals that are embedded in structures of social relationships, intentionality, and meaning. Conversely, in the nonsocial domain, emotions are likely to promote individual survival by maintaining immediate physiological and behavioral resources to biologically significant stimuli in terms of basic approach versus aversion responses (Britton et al., 2006; Insel, 2010; Porges, 2003).

Against this background, it is likely that the social versus nonsocial content of information may constitute a fundamental and distinct stimulus dimension. A number of functional magnetic resonance imaging (fMRI) studies have therefore examined the potentially different neural substrates of social versus nonsocial information processing by comparing it to the neural processing of other stimulus dimensions, particularly emotional content in terms of a positive versus negative (versus neutral) hedonic valence dissociation (Britton et al., 2006; Frewen et al., 2010; Goossens et al., 2009; Hariri et al., 2002; Norris et al., 2004; Scharpf et al., 2010; Vrtička et al., 2011, 2013). Several of these fMRI studies found brain areas showing preferential processing of social versus nonsocial information – including the occipital cortex / fusiform gyrus, amygdala, superior temporal sulcus, insula, and orbitofrontal cortex –, and consequently suggested that neural processing of the social content dimension may occur in an additive or even an interactive manner with the emotional content dimension. Only one fMRI study (Vrtička et al., 2013), however, so far directly tested whether social content and emotional content of pictorial stimuli are processed in an additive and/or interactive manner, and found that neural processing of social and emotional content interacted distinctively in bilateral amygdala, right fusiform gyrus, right anterior superior temporal gyrus, and ventromedial prefrontal cortex. In all four areas, there was a fundamental social > nonsocial activation difference for emotional (positive and negative) images, with the same effect being present for emotionally neutral images. Furthermore, a social by emotional content interaction in brain activity arose (for positive and negative stimuli): activity in response to images of social content did not significantly differ between positive and negative valence, while activity for nonsocial images displayed a negative > positive valence effect. Described in other terms, there was a significantly larger social versus nonsocial activation difference for positive as compared to negative images. Importantly, this interaction was independent of low-level stimulus properties such as spatial frequency, contrast, and luminance, as well as arousal. Together, the above findings by Vrtička et al. (2013) corroborate the notion that social content represents a fundamental and distinct stimulus dimension, and that information pertaining to the social versus nonsocial nature of stimuli is integrated with information regarding their emotional content.

In the present study, we aimed at further characterizing the interaction between social and emotional content during complex visual scene processing. More specifically, we focused on its underlying spatio-temporal pattern by means of event-related brain potentials (ERPs) while applying a very similar experimental design as implemented by Vrtička et al. (2013) using fMRI. Because of their excellent temporal resolution, ERPs provide a powerful tool to investigate the processing specificities triggered by different types of salience over time. The most prominent ERP components sensitive to emotional salience are the *Early Posterior Negativity* (EPN) and the *Late Positivity Complex* (LPC), with the latter often likewise termed as *Late Positive Potential* (LPP) (e.g. Schupp et al. 2004). The EPN, which occurs as a relative negativity over posterior electrode sites starting around 150-200 ms after stimulus onset, has been proposed to reflect enhanced sensory encoding resulting from involuntary capture of attention by various stimuli of emotional content, (e.g., Junghoefer et al., 2001; Schupp et al. 2007; Schacht & Sommer, 2009a; Bayer & Schacht, 2014). The LPC/LPP has been linked to higher-order stages of stimulus evaluation, developing around 300 ms and typically lasting for several hundred milliseconds (e.g., Schacht and Sommer 2009a). Complementing these findings, there is growing evidence indicating prioritized processing of emotionally salient stimuli to start already at early sensory stages. Several studies demonstrated the amplitudes of the visual C1 (peaking around 80 ms) and P1 (peaking around 100 ms) components to be enhanced for emotional compared to neutral stimuli (Batty & Taylor, 2003; Brosch, Sander, Pourtois, & Scherer, 2008; Holmes et al., 2009; Ortigue et al., 2004; Pourtois, Grandjean, Sander, & Vuilleumier, 2004; Stolarova, Keil, & Moratti, 2006; Rossi et al., 2017; Rellecke et al.,2012).

In contrast to the well documented ERP modulations by emotional relevance, effects of other sources of relevance, including social relevance, and their integration with emotional aspects have been largely neglected to far. Only one previous EEG study (Okruszek et al., 2016) explicitly aimed at differentiating social from nonsocial content in addition to testing for emotional content effects using complex visual scenes, although this study only comprised stimuli with a negative versus neutral valence. The authors reported early effects of social content at the P1 (social > nonsocial) and at the EPN (nonsocial > social) component. Later stages of processing were only impacted by emotional content, as indicated by larger P3 and LPP amplitudes for negative than neutral picture content. Interactions between social and emotional content were restricted to the N2 component, with extenuated amplitudes for negative pictures with social content compared to all other picture conditions. Although these findings provide first insight into the temporal dynamics of the processing of social and emotional content, they lack information on the neural processing of positive valence and are further inconclusive due to highly unconventional choice of electrodes and quantification of ERP amplitudes. Together, it remains an open question in which temporal sequence different stimulus dimensions are processed by the human brain, and what significance such processing sequence may have for physiological responses and behavior.

One theoretical framework that is devoted to addressing this question is the appraisal theory of emotion (see e.g. Sander et al., 2005; Scherer, 2009). Importantly, this theory comprises a component process model of emotion that proposes a sequence of appraisal checks that coordinate a range of responses to a particular event. Within this approach, the detection of relevance is considered to be “a first selective filter through which a stimulus or event needs to pass to merit further processing” (Scherer, 2009) (p. 3463), and to comprise information evaluation in terms of novelty (i.e. suddenness, familiarity, and/or predictability), intrinsic pleasantness (i.e. negative versus positive [versus neutral] valence), and goal / need relevance (i.e. whether the assessed information accords to or obstructs the current goals and needs of the organism).

First evidence that such temporal sequence of stimulus appraisal – particularly related to relevance detection – is implemented at the brain level was provided by an ERP study, which investigated the neural unfolding of effects of novelty and intrinsic pleasantness by means of negative, positive, and neutral images during an oddball task (van Peer et al., 2014). The authors reported a novelty effect arising first in ERPs between 200 and 300 ms, followed by an intrinsic pleasantness effect between 300 and 400 ms, and finally a novelty by intrinsic pleasantness interaction between 700 and 800 ms. The temporal dynamics of these effects indicate that the processing of intrinsic pleasantness, i.e., the emotional content in terms of positive versus negative (versus neutral) valence, is appraised relatively early during the temporal sequence and thus constitutes one of the first relevance checks, although not the very first one.

Concerning the processing of social content, no study has yet assessed its temporal unfolding over time in the context of relevance detection and the appraisal theory of emotion. As already mentioned above, however, Ondruszek et al. (2016) provided preliminary evidence for early effects of social content at the P1 (social > nonsocial) and at the EPN (nonsocial > social) component, regardless of stimulus valence. Such early social relevance effect in terms of a social versus nonsocial activation difference during complex visual scene processing is corroborated by a recent study using negative versus neutral written sentences as stimuli, and manipulating the social content dimension by means of social closeness (i.e. whether the sentences referred to participants’ significant others or to unknown agents) (Bayer et al., 2017). The authors also report early social content effects in ERPs in the P1 component (from 73 to 120 ms), and this again irrespective of the sentences’ valence. Furthermore, the authors report an effect of emotional content at a later stage in the EPN component (from around 200 ms on), and an interaction between social and emotional content in terms of the EPN having a longer duration when emotional words were presented in highly relevant social contexts, that is, referring to the participants’ boyfriend or best friend. Despite differences in the way the social content dimension was characterized, these data together point to the fact that a relevance check pertaining to social content may also occur early during stimulus appraisal – already at the P1 component –, and that an interactive processing of social and emotional content may follow at the EPN component and/or later on.

By applying a very similar experimental design as implemented by Vrtička et al. (2013) using fMRI, and according to theoretical considerations and available data on the temporal dynamics of social and emotional content processing using EEG outlined above, we predicted that social content might constitute a distinct stimulus dimension to be appraised in a relevance check separate from hedonic pleasantness. Specifically, we anticipated effects of social content to occur early during stimulus processing, presumably modulating already the P1 component of ERPs. We also predicted modulation of ERP responses by emotional content reflecting another relevance check. Although there are reports of emotional content effects occurring as early as around 100 ms after stimulus onset, the two previous ERP studies directly manipulating social and emotional content demonstrated emotion effects at subsequent ERP components, namely the N2, EPN, and P3, respectively. We therefore assumed a temporal sequence in the order of social content followed by emotional content appraisal. Finally, in line with previous fMRI data (Vrtička et al., 2013), we expected to find a robust interaction between social and emotional content according to the pattern described above that was present in bilateral amygdala, right fusiform gyrus, right anterior superior temporal gyrus, and ventromedial prefrontal cortex. Although scalp-surface ERPs cannot capture neural activity in the amygdala, we nonetheless anticipated to observe a social by emotional content interaction in cortical areas like the fusiform gyrus and/or anterior superior temporal gyrus.

## Method and Materials

### Participants

The experiment was conducted with 24 participants ranging in age between 20 and 33 years (M = 25.71 years, SD= 3.42). Only female participants were accepted for participation as emotion and social content effects may differ between sexes, with females tending to show stronger effects (Bennett et al., 2005; Federmeier et al., 2001). All participants had normal or corrected-to-normal vision and, according to the Edinburgh’s Handedness Inventory (Oldfield, 1971), were right-handed. Participation was voluntary, and the study was approved by the ethics committee of the Institute of Psychology, University of Göttingen. Participants gave written informed consent prior the study and were reimbursed for participation.

### Materials

Out of a set of 360 previously validated colored images depicting complex visual scenes (Vrtička et al., 2011, 2013), 120 pictures were chosen for the current study. All images were collected from the International Affective Picture System (IAPS) or from free sources of the Internet. They varied in their social (social, nonsocial) and emotional content (positive, negative, neutral) resulting in six experimental conditions (with 20 images per condition). All 120 images were adjusted on low-level properties, including luminance, *Fs*(1,18) < .953, *ps*> .40, contrast, *Fs*(1,18) < .37, *ps*> .69, as well as high, *Fs*(1,18) < 2.29, *ps* > .13, and low spatial frequency, *Fs*(1,18)< 3.16, *ps* > .07. Furthermore, emotional valence and arousal was controlled via pre-experimental ratings (Vrtička et al., 2011, 2013). Positive images had higher valence,*F*(1,19)= 6530, *p* < .001, and lower arousal, *F*(1,19) = 1269, *p* < .001, ratings than negative images, but there were no differences in those measures related to social content, *Fs*(1,19)< .106, *ps* > .75, and there was no interaction between emotional and social content, *Fs*(1,19)< .862, *ps*> 365. For the neutral control condition, emotional valence ratings (on a scale from 0 to 100; mean= 49.75) were situated between positive (mean= 77.80) and negative (mean= 16.15) images, and arousal ratings (mean= 33.09) were also situated between positive (mean= 43.73) and negative (mean= 73.77) images. However, emotional valence and arousal ratings did not differ between neutral social and nonsocial images (paired t-test, two tailed, *ps*> .21).

For the present experiment, scrambled versions of the 120 original images were created using Adobe Photoshop (version 11; “Filter / Telegraphics / Scramble” command), thereby generating another set of 120 images consisting of 3072 randomly distributed small squares each.

### Procedure

Before the start of the experiment, participants signed informed consent and provided demographic information. Stimuli were presented at the center of a computer screen (grey background), positioned at a distance of 90 cm from the participant. Stimuli presentation was controlled by Presentation^®^ Software.

The main experiment consisted of four blocks. Within each block, all 240 images –120 intact (target) and 120 scrambled (distractors) – were presented in randomized order. The participants’ task was to indicate by button press whether the presented image was intact or scrambled. Response-by-button assignments were counterbalanced across participants. Each trial started with the presentation of a fixation cross for 2500 ms, followed by the picture stimulus shown for 150 ms. After a blank of 850 ms duration, a question mark was presented for maximum 3000 ms, indicating the time period for responses. Feedback (“too fast”, “too slow”) was provided in case of responses outside of this interval. With the button press, the next trial was initialized. Breaks were included after every 120 trials. In order to familiarize participants with the timing of stimulus presentation and procedure of the task, there were 10 practice trials (half distractors) prior the experiment.

Picture stimuli consisted of 512 × 384 Pixel (14 × 10.5 cm), corresponding to a visual angle of 8.8° × 6.7°. Fixation crosses, feedback stimuli, and question marks were presented in white color.

### Electrophysiological Recordings and Pre-processing

The electroencephalogram (EEG) was recorded from 64 electrodes placed in an electrode cap (Easy-Cap, Biosemi, Amsterdam, Netherlands) according to the extended 10-20 system (Pivik et al., 1993). The common mode sense (CMS) electrode and the driven right leg (DRL) passive electrode were used as reference and ground electrodes (cf. www.biosemi.com/faq/cms&drl.htm). Six external electrodes were placed laterally and inferior to the eyes to record blinks and eye movements, and on the left and right mastoids. Signals were recorded at a sampling rate of 512 Hz and a bandwidth of 104 Hz and offline filtered with a Low Cutoff (0.03183099 Hz, Time constant 5 s, 12 dB/oct), a High Cutoff (40 Hz, 48 dB/oct), and a Notch Filter (50 Hz). Data was processed with BrainVision Analyzer (Brain Products GmbH, Munich, Germany). Data was average-referenced and corrected for blinks and eye movements using Surrogate Multiple Source Eye Correction (MSEC) (Ille et al., 2002) as implemented in BESA (Brain Electric Source Analysis, MEGIS Software GmbH, Gräfeling, Germany). The continuous EEG signal was segmented into epochs of 1100 ms, starting 100 ms before stimulus onset and referred to a 100 ms pre-stimulus baseline. After rejecting epochs containing artifacts (criteria: voltage steps larger than 50 μν, 200μV/200ms intervals difference of values, amplitudes exceeding -150 μV/150 μV, and activity smaller than 0.5 μV), ERP segments were averaged per participant and experimental condition.

### Data analyses

Reaction times (RTs) were analysed with a repeated-measures analysis of variance (rmANOVA), including the factors social content (social, nonsocial) and emotional content (positive, negative). In addition, we tested effects of social content in performance to neutral pictures. Accuracy was calculated as an average over all conditions and participants.

For EEG data analysis, data from practice trials, the first trial of each block, trials with erroneous or missing responses, and distractor trials (trials containing scrambled pictures) were discarded. In order to allow direct comparisons of results between the present ERP and the previous fMRI study (Vrtička et al., 2013), analyses were conducted for testing the social by emotional content interactions on positive/negative pictures and neutral pictures separately.

ERP data was analysed as follows: Based on previous research and visual data inspection, time windows for ERP components of interest were chosen as follows: i) P100 between 80 and 120 ms, ii) EPN between 200 and 320 ms, iii) P300 between 320 and 420 ms, and iv) LPC between 420 and 620 ms. The P100 component was quantified by mean amplitudes at PO7 and PO8 electrodes, where the component showed their maximal positivity. EPN amplitudes were averaged across electrodes P9, PO7, O1, Iz, Oz, O2, PO8, P10, covering typically involved posterior electrode sites (hereafter named ‘posterior ROI’). P300 and LPC mean amplitudes were first quantified at a cluster of parietal electrodes, including P3, P1, Pz, P2, P4, PO3, POz, and PO4 (‘parietal ROI’). However, as becomes visible in Figure 2B, the distribution of ERP components within the two latter time windows was shifted towards posterior sites, being highly similar to the preceding EPN time window. We therefore applied the same posterior ROI also to the ERP analyses during the P3 and LPC time intervals.

Mean amplitudes were analyzed with rmANOVAs, including the factors social content (2, social, nonsocial), emotional content (2, positive, negative), and electrode (2 or 8, respectively). In case of significant main effects or interactions between the experimental factors included, follow-up analyses were conducted with paired samples t-tests, integrated by bootstrapped (10.000 samples) 95% confidence intervals of mean differences. In line with analyses of behavioral data, ERPs to pictures of neutral content were analyzed separately. Here, the same clusters of electrodes and time windows were used as in the analyses described above.

In order to estimate the neural generators underlying the dominant voltage topographies identified at the scalp level, sLORETA (Pascual-Marqui, 2002) was used. sLORETA is a distributed linear inverse solution based on the neurophysiological assumption of coherent coactivation of neighboring cortical areas, that are known to have highly synchronized activity (Dasilva, 1991). Accordingly, it estimates multiple simultaneously active sources without any a-priori assumption on the number and position of the underlying dipoles. sLORETA solutions are computed within a three-shell spherical head model co-registered to the MNI152 template (Mazziotta et al., 2001). sLORETA estimates the 3-dimensional intracerebral current density distribution in 6239 voxels of 5 mm spatial resolution. We performed comparisons on log-transformed data using paired-samples t-tests in the time windows corresponding to relevant ERP effects. Only one single t-test per voxel was performed per time window. Statistical analyses were based on a stringent nonparametric randomization (5000 iterations), providing corrected *p*-values. Given the low resolution of sLORETA, only brain areas showing a minimum of k> 20 significant voxels will be reported, together with coordinates referring to maximum activations within brain areas.

Additional exploratory analyses were conducted to estimate the relative onsets and of effects of the two factors social and emotional content. To this aim, we calculated Global Field Power (GFP; (Lehmann and Skrandies, 1980) for all participants and experimental conditions. GFP reflects the overall ERP activity across the scalp at any given moment. Mean amplitudes of GFP were averaged in consecutive 40-ms time windows between 80 and 600 ms. In addition to this approach, main effects of experimental factors on ERPs were tested for statistical significance through two-tailed non-parametric permutation tests based on the *t*_max_ statistic (Blair and Karniski, 1993). These analyses were performed using the Mass Univariate ERP toolbox written in Matlab with a family-wise alpha level of 0.05 (Groppe et al., 2011a, b), separately for emotional content (positive vs. negative), social content (social vs. nonsocial), and for social content (social vs. nonsocial) in neutral pictures only. This approach avoids the a-priori definition of temporal and/or spatial regions of interest, since the relevant univariate test comparing participants’ ERP amplitudes in different conditions is performed for each channel-time pair. Corresponding to the combination of the 64 electrodes and 500 time points included between 0 and 1000 ms post-stimulus, 32,000 comparisons were performed in each of the three analyses. Each comparison was repeated 2,000 times. Therefore, the most extreme *t*-value (i.e., the *t*_max_) in each of the 2,000 permutations was used to estimate the *t*_max_ distribution of the null hypothesis against which to compare the 32,000 observed *t* values.

## Results

### Behavioral Performance – Reaction Times and Accuracy

Reaction times (minimum, maximum, mean, and standard deviation) for all 6 intact picture experimental conditions are summarized in Table 1 below. rmANOVAs with the factors social and emotional content for intact images did not reveal any significant main effects or interactions (all *ps* > .20). Using paired samples t-tests, two tailed, we did not find any significant differences between social and nonsocial neutral intact images, either (all *ps* > .55). Mean accuracy (percent correct responses to the task – intact versus scrambled decision) was very high at a value of 99.42 +/-1.06%.

**Table 1.** Reaction times for each intact picture experimental target condition.

|  | Minimum | Maximum | Mean | Standard Deviation |
| --- | --- | --- | --- | --- |
| *Positive Nonsocial* | 222.82 | 583.3 | 379.70 | 111.58 |
| *Positive Social* | 231.93 | 592.02 | 379.50 | 119.74 |
| *Neutral Nonsocial* | 232.46 | 561.36 | 378.98 | 103.87 |
| *Neutral Social* | 214.56 | 587.98 | 382.03 | 116.52 |
| *Negative Nonsocial* | 205.77 | 612.86 | 372.70 | 114.25 |
| *Negative Social* | 219.3 | 606.47 | 377.25 | 114.64 |

### ERP Modulation by Social and Emotional Content

*ERP Effects of Social Content.* Averaged ERPs contrasted for social versus nonsocial content are depicted in Figure 1A and B. During the P1 time window (80 to 120 ms), the rmANOVA on mean amplitudes – quantified at electrode sites PO7 and PO8 – revealed a significant main effect of social content, *F*(1,23) = 7.522, *p* = .012, *η_p^2^_* = .246, with augmented amplitudes for pictures of social content compared to pictures of nonsocial content, *T*(23) = 2.743, *p* = .020, CI(95%) = [0.153, 0.814] (Figure 1B, left panel). A similar pattern occurred during the EPN interval (200 to 320 ms) – quantified at posterior electrode sites –, by means of a significant main effect of social content, *F*(1,23) = 14.731, *p* = .001, *η_p^2^_* = .390, which was driven by larger pos terior positivities of pictures of social in comparison to nonsocial content, *T*(23) = 3.838, *p* = .001, CI(95%) = [.457, 1.387] (Figure 1B, right panel). During the P3 (320 to 420 ms) and LPC (420 and 620 ms) time windows, there was no significant main effect of social content on posterior ERP activity, all *Fs* < 2.0, *ps* > .1.

**Figure 1.**
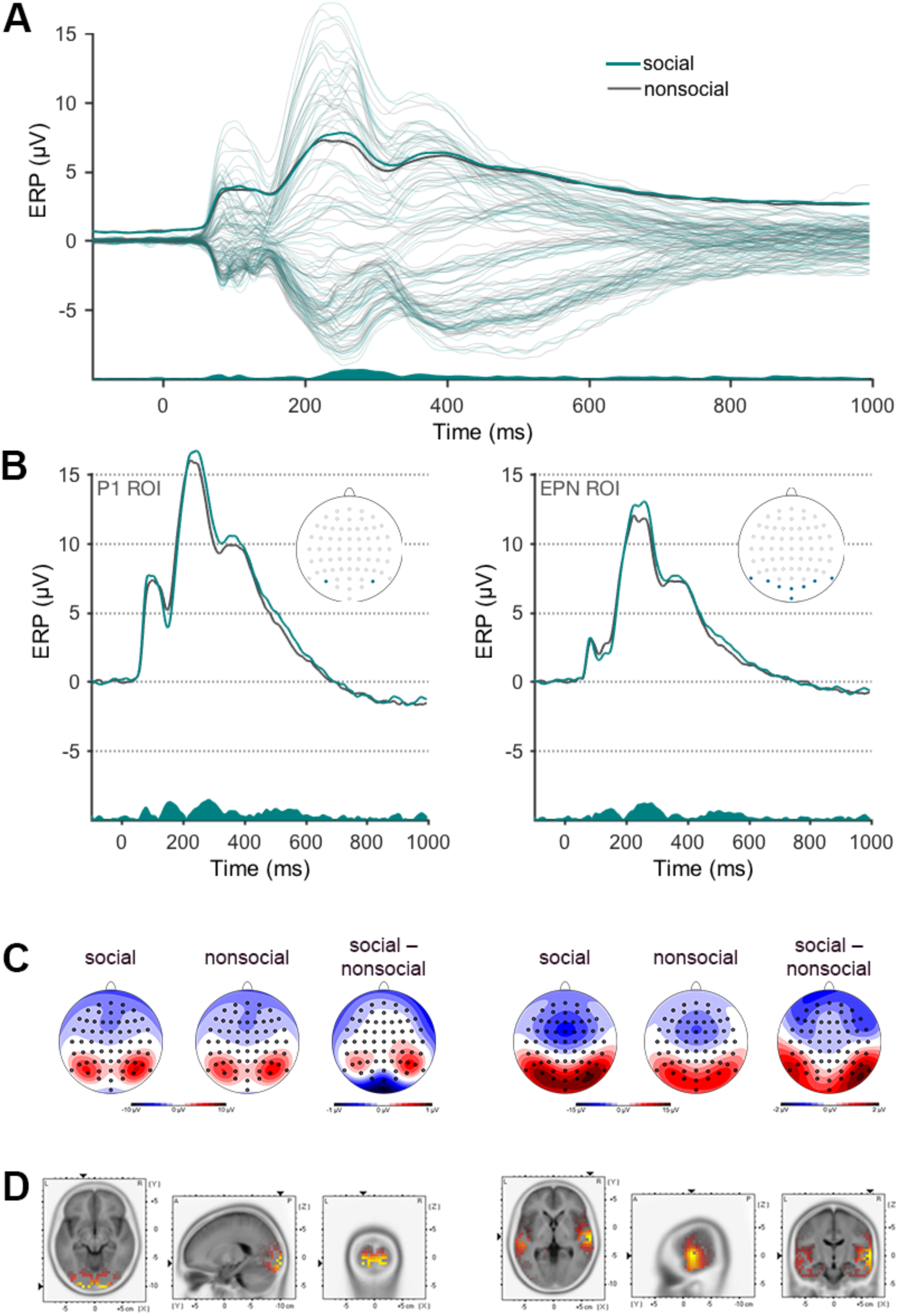
ERP effects of social content. **A** Grand average ERPs, contrasted for social and nonsocial picture content and time-locked to stimulus onsets. Signals at all EEG channels are plotted superimposed; Global Field Power (GFP) is highlighted. Inserted areas resemble the difference between the two conditions of interest over time **B** Grand average ERPs for social and nonsocial content averaged over P1- and EPN-ROI electrodes. Inserts highlight selected ROI electrodes. **C** Scalp distribution of ERP effects within the P1 (left) and EPN (right) time windows, and **D** respective source localizations of the social > nonsocial ERP differences.

As visible in Figure 1C (left panel), the scalp distribution of the early ERP modulation by social content resembled a typical P1 component with bilateral local maxima at occipital electrode sites (left panel), whereas during the subsequent time interval (EPN), a posterior positivity – instead of the expected enlarged negativity – occurred, extending to more temporal areas and accompanied by a stronger and more widely distributed frontal negativity (Figure 1C, right panel).

Source estimations for ERP activity in the two relevant time intervals confirmed this impression: While the early P1 modulation was mainly generated in occipital brain areas, sources of the subsequent ERP effect were located in more temporal brain areas (Table 2 and Figure 1D).

**Table 2.**
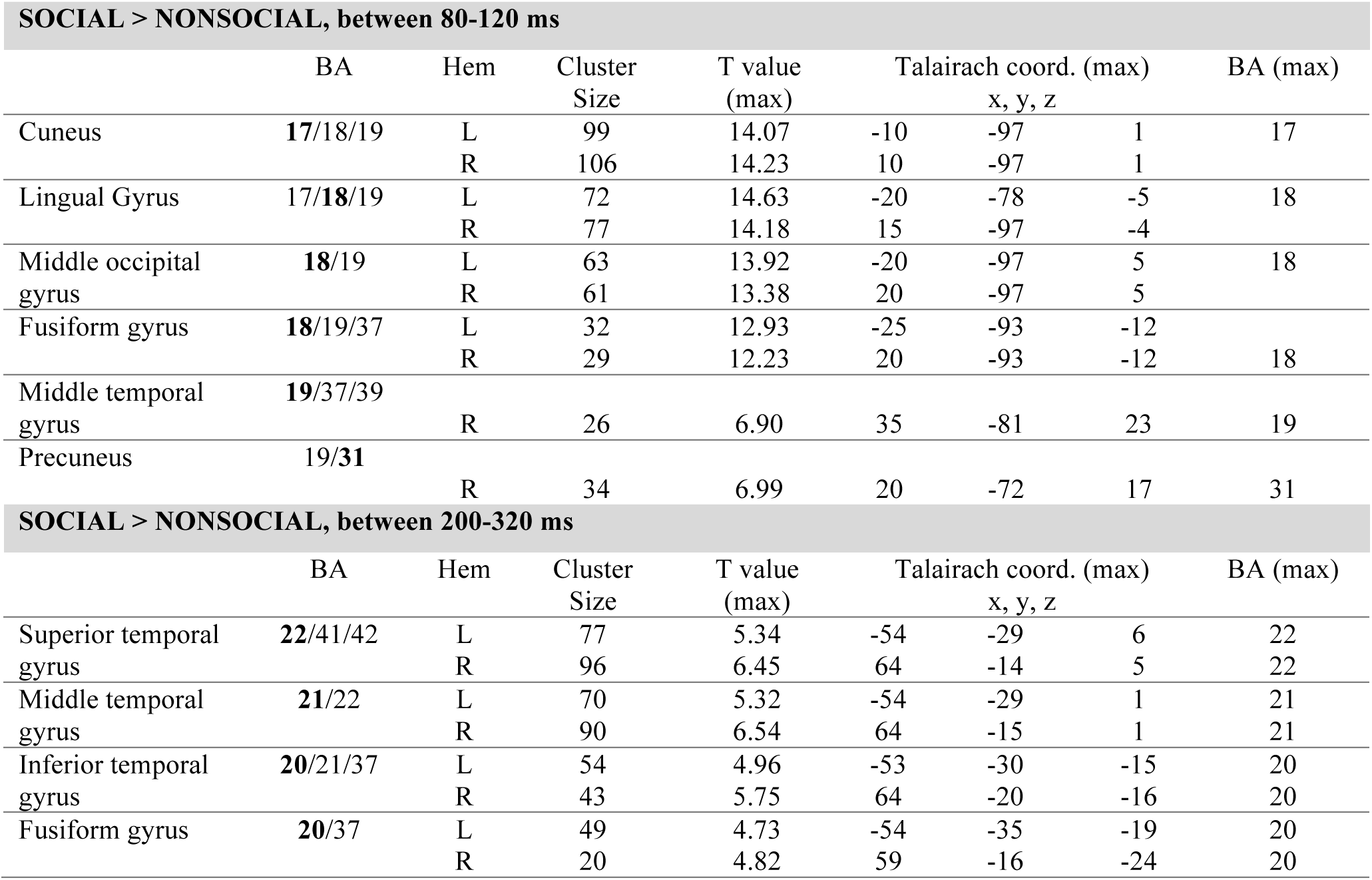
Results of source analyses of the ERP effect of social content. The list was limited to brain regions showing k > 20 significant voxels in order to account for the low resolution of the sLoreta-approach. BA= brodmann area, Hem= hemisphere.

*ERP effects of Emotional Content.* Average ERPs for negative versus positive emotional content between 200 ms and 1000 ms after stimulus onset are depicted in Figure 2A, top panel. P1 mean amplitudes – quantified at electrode sites PO7 and PO8 – were not affected by emotional content, *F*(1,23) < 1. During the EPN time window (200 to 320 ms) within the posterior ROI, however, a main effect of emotional content occurred, *F*(1,23) = 49.128, *p* = .0001, *η_p^2^_* = .681, reflecting increased posterior positivities elicited by pictures of negative valence than pictures of positive valence, *T*(23) = 7.009, *p* = .001, *CI*(95%) = [.734, 1.284]. As becomes obvious in Figure 2 (bottom panel), this posterior positivity sustained over the subsequent time intervals, i.e. between 320 and 420 ms and between 420 and 620 ms. The rmANOVAs revealed main effects of emotional content in both time intervals, *F*(1,23) = 31.118, *p* < .001, *η_p^2^_* = .575, and, *F*(1,23) = 36.731, *p* < .001, *η_p^2^_* = .611, reflecting enhanced posterior positivities to negative pictures compared to positive pictures, *T*;(23) = 5.578, *p* < .001, *CI*(95%) = [.821, 1.788], and,*T*(23) = 6.006, *p* < .001, *CI*(95%) = [.793, 1.544]. *Social by Emotional Content Interactions.* During the P1 time window (80 to 120 ms), rmANOVA on mean amplitudes at PO7/PO8 electrodes revealed a significant interaction between social and emotional relevance, *F*(1,23) = 9.910, p = .005, ηp2 = .301 (Figure 3A left panel). This interaction was driven by pictures of positive valence showing a significant social > nonsocial difference in amplitudes, *T*(23) = 3.935, *p* = .001, *CI*(95%) = [0.548, 1.556], a difference that was absent for pictures of negative valence, *T*(23) < 1 (Figure 3B, left panel).

**Figure 2.**
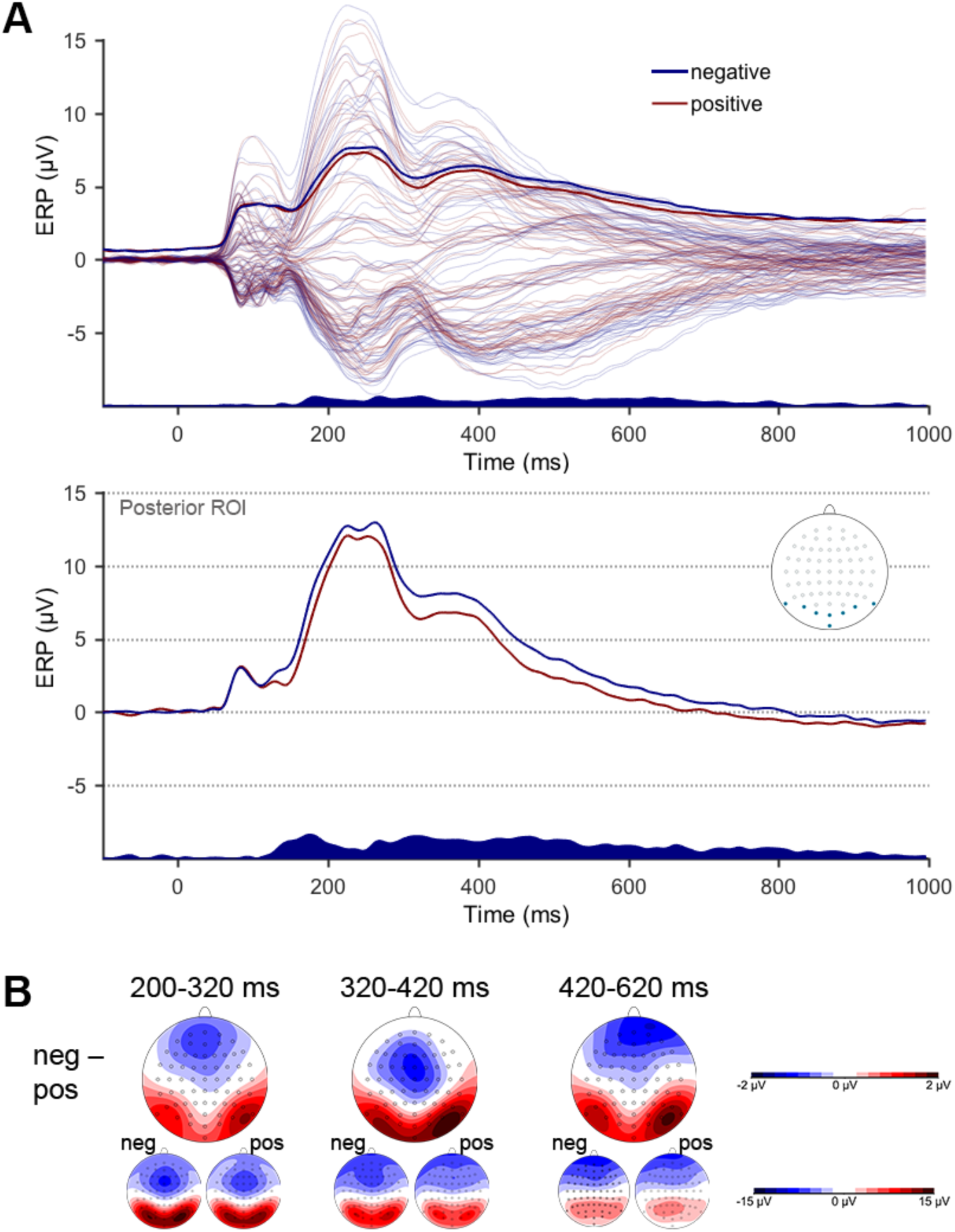
ERP effects of emotional valence. **A** Grand average ERPs, contrasted for negative and positive picture content and time-locked to stimulus onsets. Signals at all EEG channels are plotted superimposed; Global Field Power (GFP) is highlighted (upper panel). Lower panel depicts grand average ERPs averaged over posterior ROI electrodes; electrode positions are highlighted in the embedded head. Inserted areas resemble the difference between the two conditions of interest over time. Embedded head highlight selected ROI electrodes. **B** Scalp distribution of ERP effects within the three time windows of significant main effects.

During the EPN time window (200 to 320 ms) within the posterior ROI, we again observed a significant interaction between social and emotional content, *F*(1,23) = 5.256, *p* = .031, *η_p^2^_* = .186 (Figure 3, right panel). Here, however, the interaction emerged because pictures of social in comparison to nonsocial content elicited larger posterior positivities, *T*(23) = 3.838, *p* = .001, *CI*(95%) = [.457, 1.387], similar to pictures of negative compared to positive valence, *T*(23) = 7.009, *p* = .001, *CI*(95%) = [.734, 1.284]. Furthermore, differences between ERPs to negative versus positive valence were more pronounced within the nonsocial condition, *T*(23) = 5.244, *p* = .000, *CI*(95%) = [.957, 2.054], than within the social condition *T*(23) = 2.141, *p* = .045, *CI*(95%) = [.059, .958] (Figure 3B, right panel). The emotional by social content interaction did not reach significance during the P300 (320 to 420 ms) and LPC (420 and 620 ms) time intervals, all *Fs* < 2.0, *ps* > .1.

**Figure 3.**
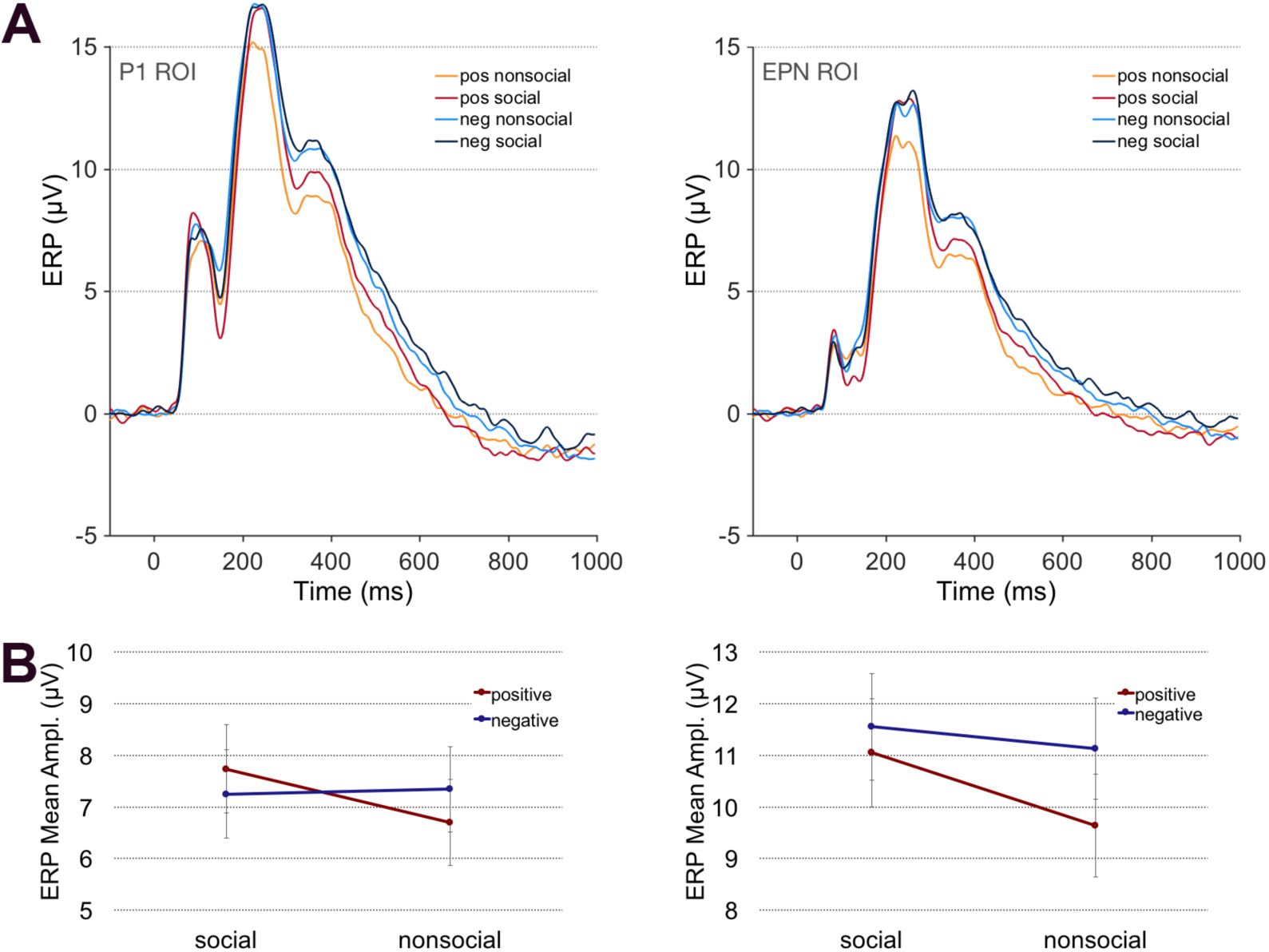
ERPs for the social by emotional content interaction. **A** Grand mean ERPs, contrasted for all conditions, averaged across P1 ROI (left panel) and EPN ROI (right panel) electrodes. **B** ERP mean amplitudes (with S.E.M.s) within the P1 time window (80-120 ms; left panel) and the EPN time window (200-320 ms).

**Figure 4.**
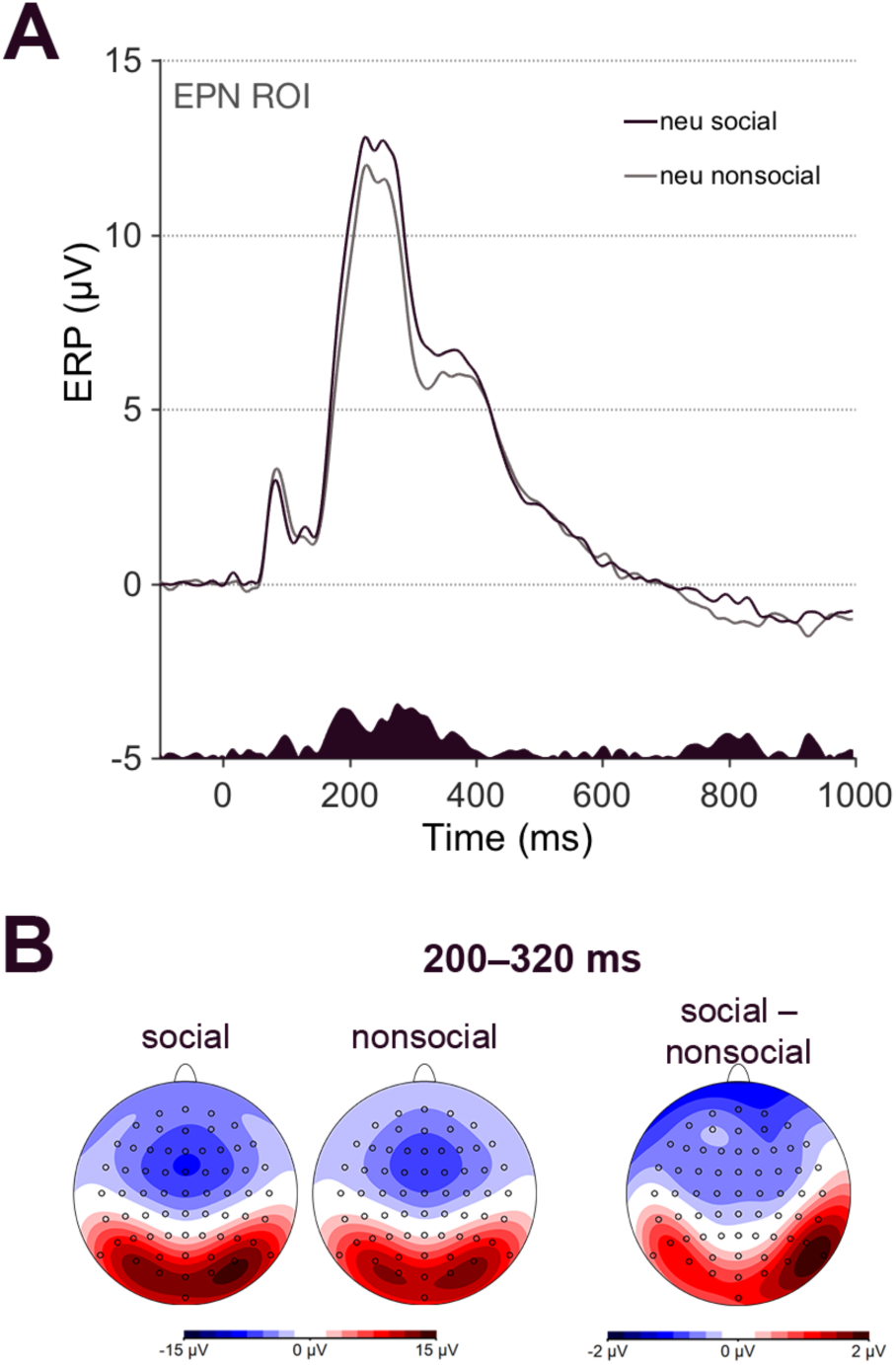
ERP effects of social relevance during neutral picture processing. **A** Grand average ERPs, contrasted for social and nonsocial content. **B** Scalp distributions of grand average EPRs and their difference between 200 and 320 ms.

### ERP components to social content in emotionally neutral pictures

In order to test potential influences of social content on the processing of emotionally neutral pictures, bootstrapped t-tests on mean ERP amplitudes were conducted. Mean amplitudes were quantified at ROI amplitudes and within time intervals indicated by significant main effects of social content in the analyses on ERPs described above.

The P1 component elicited by neutral pictures was unaffected by Social content, *T*(23) = -.632, *p* = .535, *CI*(95%) = [-0.749, 0.325]. Between 200 and 320 ms pictures of social content elicited larger posterior positivities than pictures of nonsocial content, *T*(23) = 4.653, *p* < .001, *CI*(95%) = [-0.706, 1.683], with highly similar scalp distribution as emotional pictures of social content.

### Exploratory analyses of ERPs

According to the rmANOVAs on GFP measures in consecutive 40-ms intervals, effects of Social content were restricted to the following time intervals: 80 to 120 ms, *F* =7.682, *p* < .011, *η_p^2^_* = .259, and 200 to 280 ms, *Fs* > 13.539, *ps* < .001, *η_p^2^_* > .370. Emotional Valence had long-lasting impacts on ERPs, evident between 160 ms to 720ms after stimulus onset, all *Fs* > 5.8, ps < .025, *η_p^2^_*s > .209. Interactions between social and emotional content were restricted to two time windows, first between 80 and 120 ms, *F* = 5.168, *p* = .033, *η_p^2^_* = .189, and second between 240 and 280 ms, *F* = 5.155, *p* = .033, *η_p^2^_* = .190.

## Discussion

It is generally understood that social information is of high intrinsic relevance for the human species, likely having fueled the evolution of a dedicated social brain that is nowadays investigated by the still young field of social cognitive affective neuroscience (Cacioppo and Berntson, 1992; Dunbar, 1998, 2009; Hariri et al., 2002; Keltner and Kring, 1998; Lieberman, 2007; Porges, 2003; Tomasello, 2014). Accordingly, previous functional magnetic resonance imaging (fMRI) results suggest that social content may represent a distinct stimulus dimension (Britton et al., 2006; Frewen et al., 2010; Goossens et al., 2009; Hariri et al., 2002; Norris et al., 2004; Scharpf et al., 2010; Vrtička et al., 2011, 2013). Not much is known, however, about the temporal unfolding of neural activity underlying social (versus nonsocial) information processing, and it remains largely unresolved how social content may interact with other stimulus dimensions during stimulus processing, particularly with emotional content in terms of intrinsic pleasantness / hedonic valence. Here, we extend previous EEG data (Bayer et al., 2017; Okruszek et al., 2016) by showing that social content impacts very early stages of stimulus processing, reflected in modulations of the P1 and subsequent ERP components of short latencies. Social content therefore likely represents a unique stimulus dimension that is appraised during one of the first of a series of relevance checks. In addition to a long-lasting main effect of emotional content, our findings furthermore demonstrated an interaction between social and emotional relevance at the level of the P1 and a subsequent ERP component, albeit with different interaction patterns across the two time windows. These interactions indicate that early stimulus relevance checks include both social and emotional stimulus characteristics, and that both sources of relevance of incoming information are integrated very rapidly during stimulus appraisal. Such pattern implies that social and emotional relevance is neurally appraised automatically and/or unconsciously, a notion further bolstered by the fact that neither emotional valence nor social content of pictorial stimuli were relevant for the task participants were performing during EEG data acquisition. In line with our expectations, the social by emotional content interaction pattern at the EPN component reproduced the interaction pattern previously described using the same stimuli in an fMRI study (Vrtička et al., 2013), and source estimations located such integrative processing of both stimulus dimensions at highly similar neural sites within the brain. The implications of our findings, also in relation to the appraisal theory of emotion, are outlined in more detail below.

### Effects of Social and Emotional Content in ERPs

The most interesting finding of this ERP study is an ERP modulation by both social and emotional content during complex visual scene processing that occurred as occipito-temporal positivity (and as its counterpart – a frontocentral negativity) between 200 -320 ms. This ERP effect was characterized by (i) a main effect of social content (social > nonsocial; for emotional as well as neutral images), (ii) a main effect of emotional content (negative > positive), and (iii) a social x emotional content interaction, consisting of stronger effects of social content in positive than negative images and of an enhanced difference between valence conditions in nonsocial as compared to social emotional images. Not only does this pattern strongly resemble the previously observed social by emotional content interaction effect in fMRI data using the same stimuli (Vrtička et al., 2013), but the estimated sources of the ERP effects from the current study show an intriguing overlap with the anatomical locations of the fMRI activations, particularly in the (right) FG and aSTG.

Within this latency, a relative negativity over occipito-temporal sites – associated with enhanced sensory encoding resulting from involuntary capture of attention to emotional content – has been reported across a wide range of experimental tasks and stimulus domains, including words, faces, and complex scenes (e.g., Junghoefer et al., 2001; Schupp et al. 2007; Schacht & Sommer, 2009a; Bayer & Schacht, 2014). The distribution of the ERP modulation to emotional and social relevance found in our study, does not resemble the typical EPN distribution but rather shows similarities to other N2-like effects previously been reported for increased attention allocation to emotional stimuli (e.g. Lin et al., 2008; Feng et al., 2012). Taken these findings into account, our results suggest that social content can counteract a general, and often reported (REFS), bias for negative information at early processing stages.

Interestingly, interactions between pictures’ social and emotional content became already evident in ERPs of shorter latencies (between 80 and 120 ms), namely at the P1 component. Albeit the overall interaction pattern slightly differed from that in the subsequent time window, again the positive images benefited from increased social relevance during this stage of processing. The P1 component is thought to reflect attention allocation during sensory processing of stimuli in the extrastriate visual cortex, i.e. being amplified for attended relative to unattended information (Di Russo et al., 2003; Hillyard and Anllo-Vento, 1998; Luck et al., 2000). Our findings therefore indicate a processing advantage of particularly positive social information during stimulus encoding – even in the absence of direct goal/need relevance of emotional valence and social content in terms of participants’ task instructions. Importantly, the above effects were independent of low-level stimulus properties such as luminance, contrast, and spatial frequency, and the social content effect cannot be explained by arousal, either.

The two so far available EEG studies that directly manipulated social relevance (Bayer et al., 2017; Okruszek et al., 2016) reported an effect of social content, but neither main effects of emotional valence nor a social by emotional content interaction at the P1 component. Importantly, however, both studies only included negative and neutral stimulus materials. Interestingly, other investigations on emotion processing that also included stimuli of positive valence, demonstrated amplification of the P1 amplitudes by positive emotional content, for example during word and face processing ( Bayer et al., 2012; Rellecke et al., 2011). In contrast, there are reports of early negative (versus positive and/or neutral) emotion effects on the P1 component, again during word processing (Keuper et al., 2013; Zhang et al., 2014), but also during face processing (Smith et al., 2003), as well as when participants were viewing emotional pictures from the IAPS database (Delplanque et al., 2004). In none of these previous studies, however, potential variations in stimuli’s social content have been taken into account, presumably resulting in heterogeneous findings. To fully understand what specifically determines stimulus relevance and its influence on perceptual processing, it seems indispensable to consider other content differences within and across the emotion dimension. Among them, social aspects might convey the most important information.

Besides more general aspects of early neural social and emotional content encoding discussed above, one may ask the question why particularly social positive (versus nonsocial positive) stimuli entailed early attention allocation during visual processing in our study. Positive social images used here included scenes that depicted parents interacting with their children, friends having a good time together, or happy moments in the context of romantic relationships. All of these images thus portrayed a social context of safety, security, and connectedness. In turn, nonsocial positive images showed animals, food, and appealing nature scenes (e.g. tropical beaches, sunsets, etc.). One possible mechanistic explanation of our ERP effects being mainly driven by the social positive images may thus be that this stimulus category contained a particular kind of relevance that drew early attention allocation. Such notion would accord with findings from a recent study that found increased P1 amplitudes to neutral faces previously associated with monetary rewards and thus positive motivational relevance in a social context (Hammerschmidt et al., 2017). Similarly, another study (Beckes et al., 2013) reported early attentional biases towards securely conditioned faces at the P1 component during an implicit face conditioning task, the latter effect likely representing an increase in the approach relevance of secure social bonds per se, without any added positive motivational relevance. Particularly the findings of the study by Beckes et al. (2013) would support the data obtained here, where positive social relevance was intrinsic to the depicted scenes as they were not previously associated with any rewarding value. We may therefore speculate that in our study, information pointing towards social safety and security, rather than nonsocial comfort, was particularly relevant for participants. A first relevance check during the appraisal of complex visual social emotional scenes could therefore represent a rapid assessment of information to fulfill a basic motivation to feel socially safe and secure, a computation that is not required when the processed information is nonsocial. Future investigations are needed to replicate and further characterize this early social positive effect by also taking into account the context within which information is processed, and ideally also probing for inter-individual differences that may shed more light on the source of this attentional bias.

Based on previous literature we were expecting modulation of ERPs also during later stages of affective picture processing, namely during the P3/LPC time windows (e.g. Cuthbert et al., 2000; Bayer & Schacht, 2014; Schupp et al., 2007; for a review see Olofsson et al., 2008), in particular in response to negative pictures. Indeed, a long-lasting main effect of emotional valence occurred in the present study, persisting for several hundred milliseconds, with increased ERP amplitudes to negative compared to positive images. This effect, although slightly changing in topography over time, however, did not resemble centro-parietal positivities typical for P3/LPC effects, but rather consisted of increased bilateral positivities over occipital electrode sites and – as their counterparts – fronto-central negativities. It appears difficult to compare our present data with previous studies that refrained from depicting topographical maps on the effects of interest. In a previous study that employed the same task as we used in our study (decisions on intact pictures versus their scrambled versions), grand averaged ERPs to more arousing pictures showed high similarities to our findings (Rosenkrants et al., 2008). The specific, and presumably, task-related conditions under which typical emotion-related ERPs, like the EPN and subsequent P3/LPC components are elicited, needs further investigation.

### Theoretical Implications

According to appraisal theories of emotion and the component model of emotion processing they propose (see e.g. Sander et al., 2005; Scherer, 2009), stimulus appraisal consists of a series of appraisal checks, of which the first one is devoted to relevance detection in terms of a “selective filter through which a stimulus or event needs to pass to merit further processing” (Scherer, 2009) (p. 3463). Within this framework, relevance detection is argued to comprise information evaluation in terms of novelty (i.e. suddenness, familiarity, and/or predictability), intrinsic pleasantness (i.e. negative versus positive [versus neutral] valence), and goal / need relevance (i.e. whether the assessed information accords to or obstructs the current goals and needs of the organism). First evidence on the neural temporal sequence of appraisal processes and in particular relevance detection in terms of novelty and pleasantness is already available (van Peer et al., 2014), pointing to a sequence of novelty (between 200 and 300 ms) to intrinsic pleasantness (between 300 and 400 ms) and finally a novelty by intrinsic pleasantness interaction (between 700 and 800 ms). However, such sequence originates from a particular task comprising a few novel items amongst many repeated distractors (i.e., oddball paradigm) not dissociating between stimulus dimensions in terms of social and emotional content. Using a different experimental paradigm and by directly manipulating social and emotional stimulus content, we show here that relevance detection may already occur as early as after 80 to 120 ms after stimulus onset at the P1 and continue during the EPN (between 200 and 320 ms) component, and that such relevance detection could be characterized by an interactive processing of social and emotional information. Our new findings therefore tentatively suggest that, apart from novelty, intrinsic pleasantness, and goal/need relevance, social content evaluation may represent an additional, independent early relevance check. At the same time, our data imply that different relevance checks may occur simultaneously, even already as early as 100 ms after stimulus onset, and that the outcomes of these independent relevance checks are integrated from the very beginning of stimulus appraisal. Such integrative processing of several stimulus dimensions appears to make a lot of sense particularly regarding social and emotional content of information, because differently valenced social versus nonsocial cues may signal distinct situational properties that require distinct psychological and physiological responses. In future studies, it will be important to replicate and extend the present findings by investigating even more stimulus dimensions within one experiment to capture relevance detection to novelty, intrinsic pleasantness, goal/need relevance, and social content evaluation, and particularly their inter-relations, by using different experimental paradigms in different contexts, ideally also including inter-individual differences to assess possible underlying motivational states.

### Limitations

One potential limitation of the present study is the inclusion of female participants only. It is possible that females appraise complex visual scenes differently from males in terms of particular combinations of social and emotional stimulus content possibly being attributed with a different relevance as a function of participant sex. For example, one previous EEG investigation reports sex differences in P1 amplitude modulation during emotional face viewing overall and particularly when faces were emotionally positive / rewarding (Pfabigan et al.,2014) At the same time, there is evidence for a female negativity bias at the N1 and N2 amplitudes during passive viewing of emotional images (Gardener et al., 2013; Lithari et al., 2010). Future studies are needed to resolve this issue by including both female and male participants and directly comparing brain responses between the two sexes.

Another possible limitation in the context of appraisal theories of emotion is the fact that we only assessed the two stimulus dimensions of social and emotional content, the latter representing the intrinsic pleasantness relevance check, but not other relevance checks as part of the component model of emotion processing (see above). More research is therefore clearly needed to obtain a more comprehensive image of the temporal unfolding of stimulus appraisal and relevance detection on a neural level.

## References

Adolphs, R., 2003. Cognitive neuroscience of human social behaviour. Nature Reviews Neuroscience 4, 165–178.

Aronson, E., 1980. The social animal. Palgrave Macmillan, New York.

Bayer, M., Rossi, V., Vanlessen, N., Grass, A., Schacht, A., Pourtois, G., 2017. Independent effects of motivation and spatial attention in the human visual cortex. Social Cognitive and Affective Neuroscience 12, 146–156.

Bayer, M., Sommer, W., Schacht, A., 2012. P1 and beyond: Functional separation of multiple emotion effects in word recognition. Psychophysiology 49, 959–969.

Beckes, L., Coan, J.A., Morris, J.P., 2013. Implicit conditioning of faces via the social regulation of emotion: ERP evidence of early attentional biases for security conditioned faces. Psychophysiology 50, 734–742.

Bennett, S., Farrington, D.P., Huesmann, L.R., 2005. Explaining gender differences in crime and violence: The importance of social cognitive skills. Aggression and Violent Behavior 10, 263–288.

Blair, R.C., Karniski, W., 1993. An alternative method for significane testing of wave-form difference potentials. Psychophysiology 30, 518–524.

Britton, J.C., Phan, K.L., Taylor, S.F., Welsh, R.C., Berridge, K.C., Liberzon, I., 2006. Neural correlates of social and nonsocial emotions: An fMRI study. Neuroimage 31, 397–409.

Cacioppo, J.T., Berntson, G.G., 1992. Social psychological contributions of the decade of the brain - doctrine of multilevel analysis. American Psychologist 47, 1019–1028.

Dasilva, F.L., 1991. Neural mechanisms underlying brain waves - from neural membranes to networks. Electroencephalography and Clinical Neurophysiology 79, 81–93.

Delplanque, S., Lavoie, M.E., Hot, P., Silvert, L., Sequeira, H., 2004. Modulation of cognitive processing by emotional valence studied through event-related potentials in humans. Neuroscience Letters 356, 1–4.

Di Russo, F., Martinez, A., Hillyard, S.A., 2003. Source analysis of event-related cortical activity during visuo-spatial attention. Cerebral Cortex 13, 486–499.

Dunbar, R.I.M., 1998. The social brain hypothesis. Evolutionary Anthropology 6, 178–190.

Dunbar, R.I.M., 2009. The social brain hypothesis and its implications for social evolution. Annals of Human Biology 36, 562–572.

Federmeier, K.D., Kirson, D.A., Moreno, E.M., Kutas, M., 2001. Effects of transient, mild mood states on semantic memory organization and use: an event-related potential investigation in humans. Neuroscience Letters 305, 149–152.

Feng, C., Wang, L., Liu, C., Zhu, X., Dai, R., Mai, X., Luo, Y.-L., 2012. The Time Course of the Influence of Valence and Arousal on the Implicit Processing of Affective Pictures. PLoS ONE 7(1): e29668.

Frewen, P.A., Dozois, D.J.A., Neufeld, R.W.J., Densmore, M., Stevens, T.K., Lanius, R.A., 2010. Social Emotions and Emotional Valence During Imagery in Women With PTSD: Affective and Neural Correlates. Psychological Trauma-Theory Research Practice and Policy 2, 145–157.

Gardener, E.K.T., Carr, A.R., MacGregor, A., Felmingham, K.L., 2013. Sex Differences and Emotion Regulation: An Event-Related Potential Study. Plos One 8.

Goossens, L., Kukolja, J., Onur, O.A., Fink, G.R., Maier, W., Griez, E., Schruers, K., Hurlemann, R., 2009. Selective Processing of Social Stimuli in the Superficial Amygdala. Human Brain Mapping 30, 3332–3338.

Groppe, D.M., Urbach, T.P., Kutas, M., 2011a. Mass univariate analysis of event-related brain potentials/fields I: A critical tutorial review. Psychophysiology 48, 1711–1725.

Groppe, D.M., Urbach, T.P., Kutas, M., 2011b. Mass univariate analysis of event-related brain potentials/fields II: Simulation studies. Psychophysiology 48, 1726–1737.

Hammerschmidt, W., Sennhenn-Reulen, H., Schacht, A., 2017. Associated motivational salience impacts early sensory processing of human faces. Neuroimage 156, 466–474.

Hariri, A.R., Tessitore, A., Mattay, V.S., Fera, F., Weinberger, D.R., 2002. The amygdala response to emotional stimuli: A comparison of faces and scenes. Neuroimage 17, 317–323.

Hillyard, S.A., Anllo-Vento, L., 1998. Event-related brain potentials in the study of visual selective attention. Proceedings of the National Academy of Sciences of the United States of America 95, 781–787.

Ille, N., Berg, P., Scherg, M., 2002. Artifact correction of the ongoing EEG using spatial filters based on artifact and brain signal topographies. Journal of Clinical Neurophysiology 19, 113–124.

Insel, T.R., 2010. The Challenge of Translation in Social Neuroscience: A Review of Oxytocin, Vasopressin, and Affiliative Behavior. Neuron 65, 768–779.

Keltner, D., Kring, A.M., 1998. Emotion, Social Function, and Psychopathology. Review of General Psychology 2, 320–342.

Keuper, K., Zwitserlood, P., Rehbein, M.A., Eden, A.S., Laeger, I., Junghoefer, M., Zwanzger, P., Dobel, C., 2013. Early Prefrontal Brain Responses to the Hedonic Quality of Emotional Words - A Simultaneous EEG and MEG Study. Plos One 8.

Lehmann, D., Skrandies, W., 1980. Rerfrence-free identification of components of checkerboard-evoked multichannel potential fields. Electroencephalography and Clinical Neurophysiology 48, 609–621.

Lieberman, M.D., 2007. Social cognitive neuroscience: A review of core processes. Annual Review of Psychology, pp. 259–289.

Lin, H., Jin, H., Liang, J., Yin, R., Liu, T., Wang, Y., 2015. Effects of Uncertainty on ERPs to Emotional Pictures Depend on Emotional Valence. Frontiers in Psychology, 6:1927.

Lithari, C., Frantzidis, C.A., Papadelis, C., Vivas, A.B., Klados, M.A., Kourtidou-Papadeli, C., Pappas, C., Ioannides, A.A., Bamidis, P.D., 2010. Are Females More Responsive to Emotional Stimuli? A Neurophysiological Study Across Arousal and Valence Dimensions. Brain Topography 23, 27–40.

Luck, S.J., Woodman, G.F., Vogel, E.K., 2000. Event-related potential studies of attention. Trends in Cognitive Sciences 4, 432–440.

Mazziotta, J., Toga, A., Evans, A., Fox, P., Lancaster, J., Zilles, K., Woods, R., Paus, T., Simpson, G., Pike, B., Holmes, C., Collins, L., Thompson, P., MacDonald, D., Iacoboni, M., Schormann, T., Amunts, K., Palomero-Gallagher, N., Geyer, S., Parsons, L., Narr, K., Kabani, N., Le Goualher, G., Boomsma, D., Cannon, T., Kawashima, R., Mazoyer, B., 2001. A probabilistic atlas and reference system for the human brain: International Consortium for Brain Mapping (ICBM). Philosophical Transactions of the Royal Society B-Biological Sciences 356, 1293–1322.

Münte, T.F., Urbach, T.P., Düzel, E., Kutas, M., 2000. Event-related brain potentials in the study of human cognition and neuropsychology. In: Grafman, J., Rizzolati, G. (Eds.), Handbook of Neuropsychology. Elsevier, Amsterdam.

Norris, C.J., Chen, E.E., Zhu, D.C., Small, S.L., Cacioppo, J.T., 2004. The interaction of social and emotional processes in the brain. Journal of Cognitive Neuroscience 16, 1818–1829.

Okruszek, L., Wichniak, A., Jarkiewicz, M., Schudy, A., Gola, M., Jednorog, K., Marchewka, A., Lojek, E., 2016. Social and nonsocial affective processing in schizophrenia - An ERP study. International Journal of Psychophysiology 107, 54–62.

Oldfield, R.C., 1971. The assessmentr and analysis of handedness: The endinburg inventory. Neuropsychologia 9, 97–113.

Pascual-Marqui, R.D., 2002. Standardized low-resolution brain electromagnetic tomography (sLORETA): Technical details. Methods and Findings in Experimental and Clinical Pharmacology 24, 5–12.

Patel, S.H., Azzam, P.N., 2005. Characterization of N200 and P300: selected studies of the Event-Related Potential. International Journal of Medical Sciences 2, 147–154.

Pfabigan, D.M., Lamplmayr-Kragl, E., Pintzinger, N.M., Sailer, U., Tran, U.S., 2014. Sex differences in event-related potentials and attentional biases to emotional facial stimuli. Frontiers in Psychology 5.

Pivik, R.T., Broughton, R.J., Coppola, R., Davidson, R.J., Fox, N., Nuwer, M.R., 1993. Guidelines for the recording and quantitative-analysis if electroencephalographic activity in research contexts. Psychophysiology 30, 547–558.

Porges, S.W., 2003. Social engagement and attachment - A phylogenetic perspective. In: King, J.A., Ferris, C.F., Lederhendler, II (Eds.), Roots of Mental Illness in Children, pp. 31–47.

Sander, D., Grandjean, D., Scherer, K.R., 2005. A systems approach to appraisal mechanisms in emotion. Neural Networks 18, 317–352.

Scharpf, K.R., Wendt, J., Lotze, M., Hamm, A.O., 2010. The brain's relevance detection network operates independently of stimulus modality. Behavioural Brain Research 210, 16–23.

Scherer, K.R., 2009. Emotions are emergent processes: they require a dynamic computational architecture. Philosophical Transactions of the Royal Society B-Biological Sciences 364, 3459–3474.

Smith, N.K., Cacioppo, J.T., Larsen, J.T., Chartrand, T.L., 2003. May I have your attention, please: Electrocortical responses to positive and negative stimuli. Neuropsychologia 41, 171–183.

Tomasello, M., 2014. The ultra-social animal. European Journal of Social Psychology 44, 187–194.

van Peer, J.M., Grandjean, D., Scherer, K.R., 2014. Sequential Unfolding of Appraisals: EEG Evidence for the Interaction of Novelty and Pleasantness. Emotion 14, 51–63.

Vrtička, P., Sander, D., Vuilleumier, P., 2011. Effects of emotion regulation strategy on brain responses to the valence and social content of visual scenes. Neuropsychologia 49, 1067–1082.

Vrtička, P., Sander, D., Vuilleumier, P., 2013. Lateralized interactive social content and valence processing within the human amygdala. Frontiers in Human Neuroscience 6.

Zhang, D., He, W., Wang, T., Luo, W., Zhu, X., Gu, R., Li, H., Luo, Y.-j., 2014. Three stages of emotional word processing: an ERP study with rapid serial visual presentation. Social Cognitive and Affective Neuroscience 9, 1897–1903.

